# Enteroscape: An agent-based model for simulating microbial dynamics and host responses in the gut ecosystem

**DOI:** 10.64898/2026.01.27.701954

**Authors:** David J. Datta, Reeta P. Rao, Elizabeth F. Ryder

## Abstract

Understanding microbial interactions within the gut microbiome is challenging due to the sheer number of microorganisms present. Each additional microbe introduces new layers of complexity, especially when studying the conditions and dynamics of infectious progression. To address this challenge, we have developed Enteroscape: an *in silico* agent-based model, grounded in previous research, laboratory experiments, and *in vivo* assays. Enteroscape enables simulation of diverse microbial interactions within a nematode host environment. In this study, we used Enteroscape to model the progression of pathogenic yeast infection and evaluate the effects of probiotic yeast treatment in the *Caenorhabditis elegans* intestine. Drawing upon empirical and literature-based behavioral rules for individual microbes, Enteroscape simulations produced emergent behaviors of the microbial population that mirror experimentally observed infection dynamics, including visual patterns of infectious progression, as well as the effect of probiotic treatment on the host lifespan. Enteroscape demonstrates how computational models can generate testable predictions that deepen our insight into microbial community interactions and their impact on host biology.

**Author Summary:** The human microbiome is a complicated ecosystem, consisting of trillions of microbes of hundreds of different species that live in the human intestine. Understanding how different species interact to maintain a healthy balanced microbiological ecosystem that keeps pathogenic microbes at bay is critically important to human health. We have developed a simple experimental system that allows us to examine the interactions of just a few thousand microbes at a time in a living host, the microscopic nematode worm, *C. elegans*. In prior laboratory work, we have demonstrated how ingestion by the worm of the human pathogenic microbe *C. albicans* shortens the worm’s lifespan, while co-ingestion of various probiotic microbial species restores a longer lifespan. In this paper, we report the creation of a computational model of this system, which we named Enteroscape. We show that Enteroscape mimics the results of the experimental system, including the visual progression of infection by a pathogenic species, the effects on worm lifespan, and the amelioration of the pathogens’ effects by a probiotic species. Enteroscape will allow us to develop and test hypotheses about the mechanisms underlying remediation by probiotic microbes, with potential applications to better probiotic treatment of human infections.

## Introduction

Within the human intestinal microbiome, tens of trillions of microbes, representing hundreds of species, thrive and propagate, forming a complex and dynamic ecosystem. The microbiota play a critical, though not yet fully defined, role in influencing the health of their host both within and beyond the gut [1]. While host - microbe interactions are important, microbe - microbe interactions are equally essential to understanding microbiome function.

Among the many reasons to investigate these dynamics, two stand out for their urgent clinical relevance: the emergence of drug resistance and opportunistic pathogens [2,3]. Drug resistance, particularly when it occurs across multiple species, has been difficult to elucidate and has become increasingly prevalent in recent decades [2]. Similarly, opportunistic pathogens often emerge from specific microbial interactions and environmental growth conditions, yet our understanding of their behavior remains limited. Most alarming is the intersection of these issues: drug-resistant opportunistic pathogens are becoming more common, prompting the Centers for Disease Control and Prevention (CDC) to identify and prioritize microbial threats with the greatest public health impact [4,5]. Notably, pathogenic fungi of the *Candida* genus are included on this list, designated as urgent or serious threats requiring aggressive and sustained action [6].

*Candida albicans* exemplifies this challenge. Although it typically exists in a benign yeast state within the microbiomes of humans and other animals, *C. albicans* can transition into a pathogenic filamentous state, forming hyphae and biofilms that damage host tissues. In this form, infections can range from mild skin rashes to severe gastrointestinal disruptions and deadly bloodstream infections [7]. Treatment options are limited, and drug-resistant strains of *C. albicans* have emerged in recent years [8,9]. Given the widespread presence of such microbes in the human body, the ability to identify what triggers transitions to pathogenicity and how other microbial or host factors modulate this process would be a powerful tool for both medicine and research. To that end, we developed Enteroscape, a computational simulation designed to investigate microbial interactions in the context of a host. By creating a digital representation of a host intestinal tract, Enteroscape leverages computational power to replicate simple biological behaviors *in silico* in order to elucidate and/or validate more complex community dynamics *in vivo*.

Enteroscape builds on previous studies of both *C. albicans* pathogenicity and the ability of probiotic yeasts to inhibit *Candida* infection [10–12]. *Candida* infection typically progresses through several stages: adhesion to host epithelial cells, filamentation, biofilm formation, and eventually host tissue damage [13]. Some species of yeast, such as *S. cerevisiae* and *I. occidentalis*, interfere with the first step of adhesion by secreting secondary metabolites, specifically aromatic alcohols like tryptophol and phenylethanol, which impair *Candida* virulence [10,11]. Higher concentrations of these molecules correlate with reduced adhesion and filamentous growth as well as decreased biofilm formation [11,12,14].

Enteroscape draws inspiration from experiments designed to evaluate probiotic treatments using *C. elegans* as an animal host [10–12,15,16]. *C. elegans* is a well-established model organism for gut microbiome studies due to its conserved innate immune system and intestinal epithelium, which mimics mammalian intestine with microvilli and mucosal secretion [16–20]. The worm’s transparent body enables direct visual observation, and its gut microbiome can be controlled by simply feeding it selected microbes [16–20]. In laboratory experiments, ingestion of *C. albicans* results in microbial accumulation and distension of the gut over the course of several days, which ultimately shortens the lifespan of the animal [11,12]. However, when probiotic yeast is introduced along with *C. albicans*, host survival increases significantly. Using these data and observations as a foundation, we constructed an *in silico* model that can simulate the same types of microbial inoculations as seen in laboratory experiments.

Enteroscape was developed to model host-microbe and microbe-microbe interactions within the *C. elegans* intestinal tract using an agent-based modeling (ABM) framework built in Netlogo [21], applying a pattern-oriented modeling approach [22,23]. ABM models distinct agent types or ‘breeds’, such as different types of microbes, defined by specific traits and rules based on experimental evidence or hypothesized behaviors [e.g. 24–27]. These agents populate a digital ‘world,’ allowing the researcher to observe emergent behavior that arises from interactions among these agents or between the agents and their ‘world’ itself. If the simulated outcomes resemble real-world patterns, this suggests that a model’s assumptions are realistic enough for its purpose [28]. The more patterns a model can accurately replicate, the stronger the evidence supporting the validity of its agents’ behaviors, thus enabling predictive simulations and hypothesis generation that can be tested experimentally, further strengthening this evidence. A computational simulation of agent interactions based on real *in vivo* data takes a step towards not only understanding the observed behavior of the organisms but also introduces the possibility of observing behavior *in silico* that may be difficult to observe *in vivo* or *in vitro*.

A major advantage of using an ABM modeling approach is that it allows individual microbial agents to have different properties or to be in different states, and to interact with each other locally in heterogeneous microenvironments, as they do *in vivo* in microbial communities [29–31]. Because the biological system Enteroscape simulates involves relatively few microbes (on the order of thousands rather than trillions), we were not concerned about computational expense that can be a problem in using models involving large numbers of agents. While many microbiome models have been developed that model bacterial metabolism extensively [for excellent reviews, see 32, 33], we made the conscious decision not to include overall microbial metabolism in Enteroscape and to focus instead on individual choices involved in biofilm formation during pathogenic infection.

Enteroscape is composed of three core parts: the host *C. elegans*, modeled as the ‘world,’ and two microbial agent types, pathogenic and probiotic yeast, which follow experimentally-based rules. Microbial agents in the model respond to conditions in the local environment to make individual choices about whether to transition from planktonic to adherent to biofilm-producing states.

Pathogenic yeast agents adhere to the intestinal lumen and form biofilms, while probiotic yeast agents secrete small molecules that prevent adhesion of pathogenic yeast. Host intestinal distention is modeled as a function of microbial accumulation and biofilm formation.

Simulation runs show that infections by pathogenic yeast agents produce intestinal distention patterns consistent with *in vivo* imaging, particularly at the anterior gut adjacent to the pharyngeal-intestinal valve. As in experimental studies, the presence of probiotic yeast agents reduces intestinal distention and prolongs host survival. Overall, Enteroscape demonstrates the feasibility and utility of a computational model to investigate host-microbe and microbe-microbe interactions. It provides valuable insights and a basis for further research directions.

## Results

We developed Enteroscape as an *in silico* model to explore enteric interactions based on experimental evidence from previous research [10–12], using a pattern-oriented approach [22,23] for parameter calibration. To demonstrate Enteroscape’s ability to simulate laboratory observations of *Candida* infections, we constructed a simplified representation of the experimental system in Netlogo. With this model, two pre-programmed scenarios were constructed within Enteroscape based on the experimental setup. One scenario focuses solely on the infectious progression of the pathogenic yeast absent of the probiotic microbes, described as the untreated scenario. The other simulates a co-inoculation treatment of the host with both pathogenic and probiotic yeast agents, described as the treated scenario. Enteroscape’s performance within these two scenarios was measured against similar *in vivo* results both visually and statistically. Here we provide both an overview of the design of the model, as well as the use of the model to simulate *in vivo* experiments. Computational details of the model are described in Methods. A complete description of the model following the Overview, Design concepts and Details (ODD) protocol [34–36] is provided as Supporting Information (S1 Appendix).

### The Enteroscape Model

#### Basic model design and interface

The Enteroscape model consists of two types of Netlogo agents: patches and turtles. Turtles act as mobile agents, with one ‘breed’ representing the pathogenic yeast and another representing the probiotic yeast. Patches are stationary agents, here used to represent the host *C. elegans*. The interface represents the anterior region of the *C. elegans* intestine (Fig. 1). Each patch corresponds to a 4-micron square, while microbial agents are modeled as spheres approximately 3 microns in diameter. Temporal progression in Enteroscape is measured in ticks, with one tick allowing one complete execution of the main ‘go’ function. On average, in the absence of biofilm build-up, it takes a microbial agent about 200 ticks to traverse the length of the simulated intestinal tract in Enteroscape (about 160 microns, approximately 20% of the length of the complete *C elegans* intestine).

**Fig 1:**
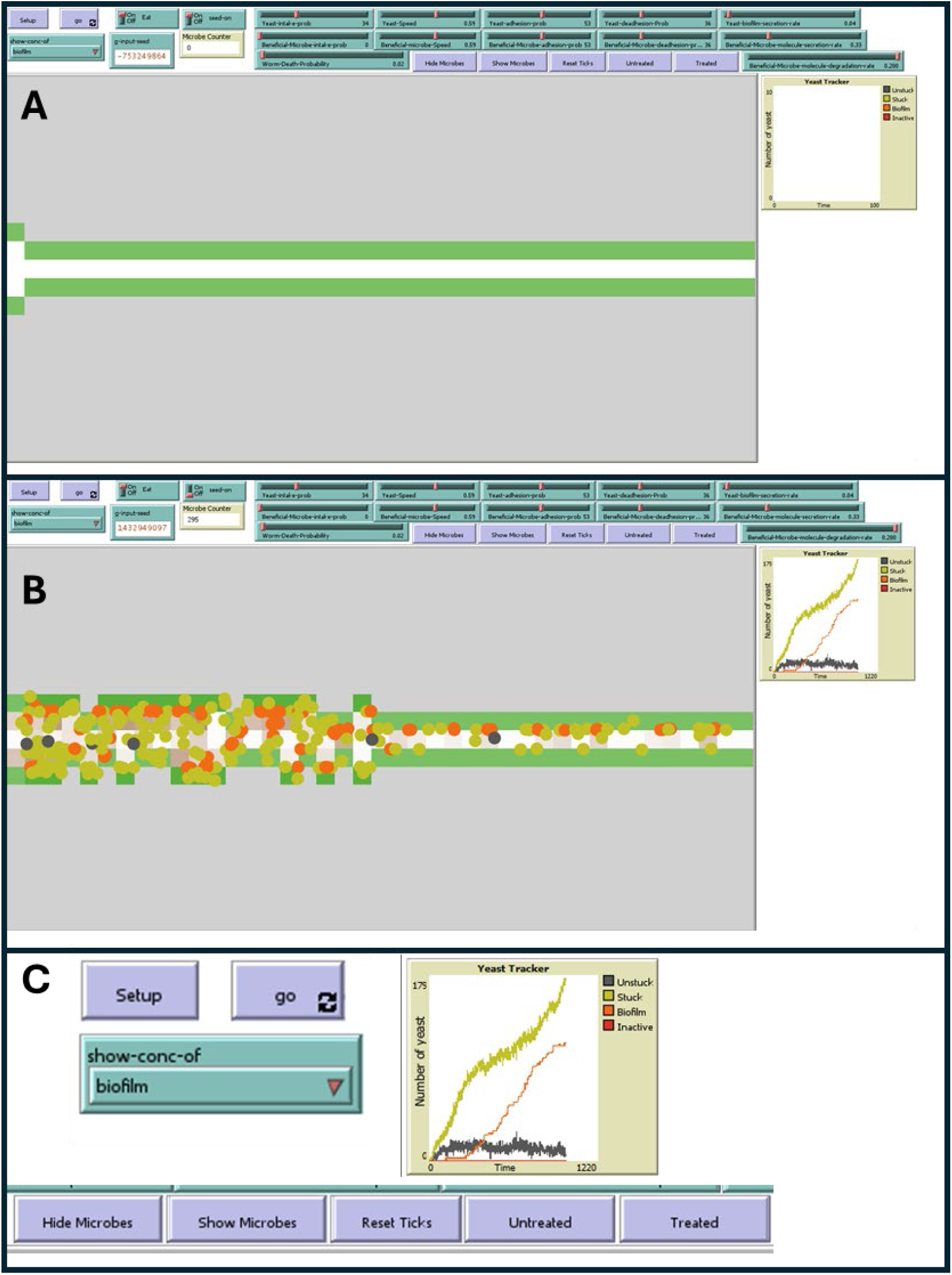
The Enteroscape interface. (A) Screenshot of initial setup of simulation. Netlogo makes use of entities called “patches” to model the *C. elegans* intestine: a grid of immovable agents, each with its own variable values and properties. Here, white patches represent the intestinal lumen, green patches represent the mucosal lining, and grey patches represent the epithelial cells of the intestine. The intestine is modeled beginning just posterior to the worm pharynx; anterior is to the left. (B) Screenshot from a simulation run including only pathogenic microbe agents. When the simulation is run, microbes are ‘ingested’ from the left of the interface (anterior of the worm). (freely moving microbes, grey circles; adhered microbes, yellow-green circles). Pathogenic microbes in biofilm state (orange) secrete biofilm into the intestine (brown patches). (C) Enlargement of some Enteroscape user input and output widgets. Enteroscape uses many of the sliders, switches, drop downs, and buttons included with Netlogo for user input on certain variables for modifying the properties of microbial agents before or during simulation. In addition to these slider variables, several tools for analyzing simulations are also provided: options for displaying gradients of small molecule or biofilm concentrations, buttons for the untreated and treated scenarios mirroring the original *in vivo* assays, a counter and graph for tracking pathogenic yeast numbers across states, and the ability to hide/show microbial agents for better visualization of secretion or distension patterning.

#### Host intestine modeling

Three types of patch agents are used for modeling the host intestine: Cell, Lumen, and Mucous (Fig. 1). The Cell patches (grey) represent the epithelial cells of the nematode intestine. During pathogenic yeast infection and subsequent distension of the intestine, Cell patches are reduced in number, indicating damage to the host. Lumen patches (white) represent the intestinal space that ingested microbes traverse. Lumen patches display a color gradient representing the concentration of either biofilm (brown) or small beneficial molecule (blue) secreted onto the patch by pathogenic or probiotic microbes, respectively. Mucous patches (green) represent the mucosal lining of the nematode intestine, and are often the initial sites of adherence for microbial agents.

#### Modeling of probiotic and pathogenic yeast

Enteroscape represents probiotic and pathogenic yeast as distinct “breeds” of agent-based entities. Although both breeds begin with the same initial behavioral states, pathogenic yeast agents include additional states that simulate their transition into pathogenic growth and biofilm formation. The modeled microbial states are summarized in Table 1, with Fig 2 describing the life cycle of pathogenic yeast agents. Probiotic yeast agents exist in either Planktonic (freely moving, light blue) or Adherent (dark blue) states (Table 1); pathogenic yeast agents can also exist as Planktonic (grey) or Adherent (yellow-green), as well as in two additional states.

**Fig 2:**
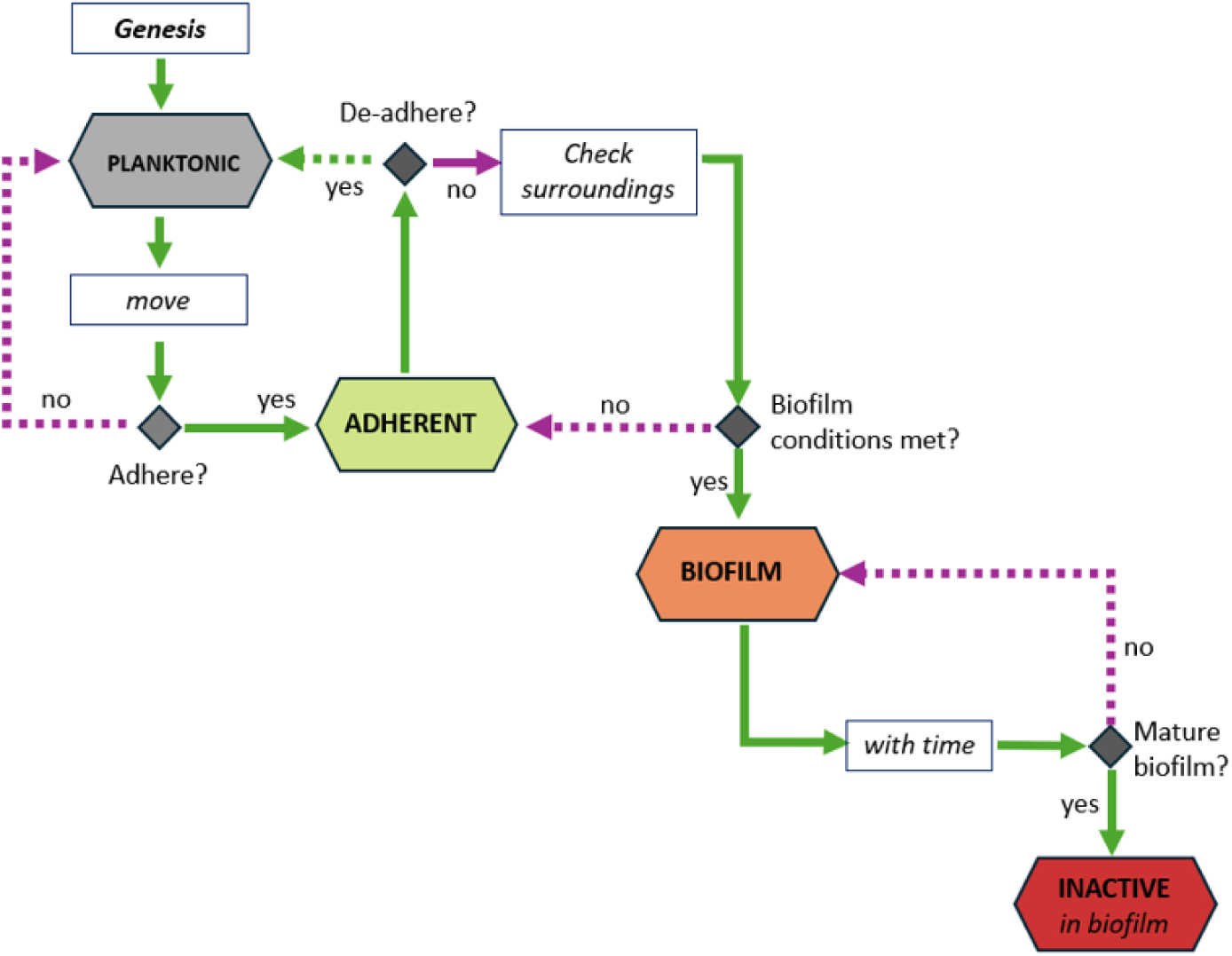
The life cycle of pathogenic yeast agents in Enteroscape. Each state (hexagon) of the pathogenic type has its own primary behaviors and functions that dictate transitions to another state. Decision points are shown as diamonds; green arrows show the response to a ‘yes’ decision, while purple arrows show the response to a ‘no’ decision.. Solid arrows indicate choices that result in the microbes transitioning to a more adherent or pathogenic state, while dotted arrows indicate choices that result in microbes transitioning to a less adherent or pathogenic state. . Each agent can make at most one state transition for each tick of the model. Starting from the genesis of the agent, this chart displays the corresponding decision flows made by agents in each state. Starting from the Planktonic state (gray), an agent will attempt to move before checking if the agent should transition to the Adherent state (yellow-green). Agents in the Adherent state check to see if the agent should revert to Planktonic state. If not, Adherent agents check their surroundings to see if they should transition to a Biofilm state (orange). Once an agent is in the Biofilm state, it cannot convert backward toward the Planktonic state; it can only move forward over time to a Metabolically Inactive state (red).

**Table 1:**
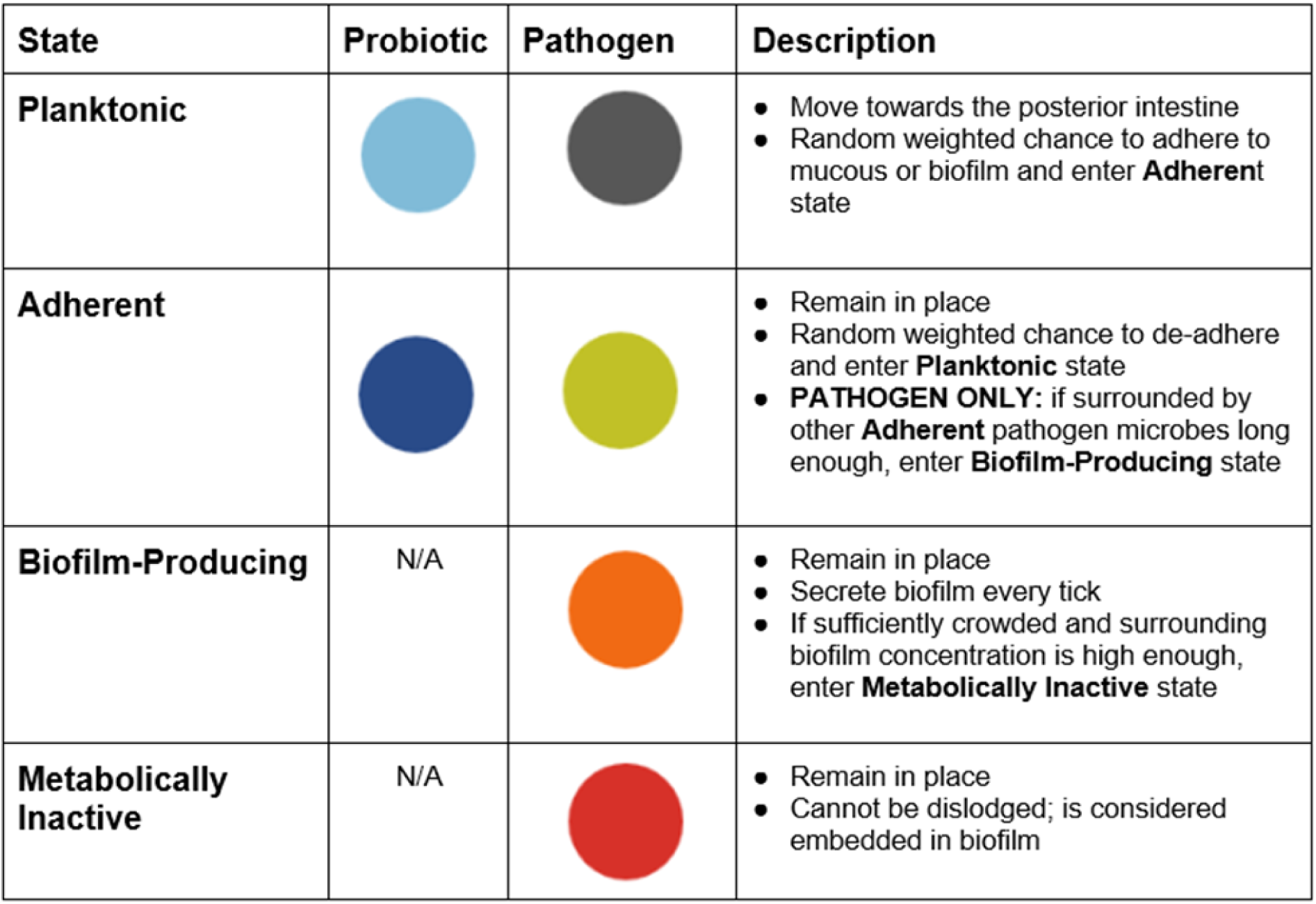
Microbial Agent states. Both pathogenic and probiotic microbes share the Planktonic and Adherent states, since they are functionally the same for both. However, the probiotic type, regardless of state, secretes small beneficial molecule constantly, while the pathogenic agent type does not secrete anything while in the Planktonic or Adherent states. For the pathogenic agent type, two additional states beyond the Adherent state exist: Biofilm-Producing and Metabolically Inactive. The Biofilm-Producing state is the only state where the pathogenic agent type secretes biofilm onto the patch it occupies and neighboring patches. Rarely, if there is enough space, these Biofilm-Producing yeast may also skim off a budding biofilm segment and relocate down the intestine. Once surrounded by high concentrations of biofilm however, Biofilm-Producing yeast will transition to the Metabolically Inactive state where they cease secreting biofilm and remain stagnant.

| State | Probiotic | Pathogen | Description |
| --- | --- | --- | --- |
| **Planktonic** |  |  | • Move towards the posterior intestine<br>• Random weighted chance to adhere to mucous or biofilm and enter **Adherent** state |
| **Adherent** |  |  | • Remain in place<br>• Random weighted chance to de-adhere and enter **Planktonic** state<br>• **PATHOGEN ONLY**: if surrounded by other **Adherent** pathogen microbes long enough, enter **Biofilm-Producing** state |
| **Biofilm-Producing** | N/A |  | • Remain in place<br>• Secrete biofilm every tick<br>• If sufficiently crowded and surrounding biofilm concentration is high enough, enter **Metabolically Inactive** state |
| **Metabolically Inactive** | N/A |  | • Remain in place<br>• Cannot be dislodged; is considered embedded in biofilm |

Probiotic agents are assumed to secrete small molecules constantly, regardless of their state; these small molecules lower the probability of adhesion of both types of microbes. In contrast, pathogenic yeast must enter the specific Biofilm-producing state (orange) in order to build a biofilm with their associated secretions (Fig. 2). If Biofilm-producing yeast become embedded deeply and long enough, they will transition to the Metabolically Inactive state (red) to mimic the hibernative state exhibited by many organisms within areas of a biofilm with low nutrient availability [14].

The behaviors and steps within this life cycle are informed by experimental evidence from the literature [10–12,16] and ongoing studies. Because complete biofilm maturation and widespread microbe proliferation have not yet been conclusively demonstrated *in vivo* within the *C. elegans* gut, Enteroscape focuses on modeling the early to mid-stages of biofilm development only. This approach allows for realistic simulation of observed behaviors without overextending assumptions beyond the available empirical data.

#### Modeling intestinal distension

In laboratory research, infection of *C. elegans* with *C. albicans* led to initial swelling of the anterior intestine that progressed towards the posterior over time [11]. This distension damaged the host, resulting in the death of the nematode as severity increased. Within Enteroscape, distension results from the accumulation of yeast agents and biofilm secretion as infection progresses in the simulation. As more pathogenic yeast agents adhere to a region, more biofilm will accumulate; once a threshold is reached, the intestine will ‘swell’; Mucus cells appear to move outward from the intestinal center to mimic *in vivo* observations. The threshold at which distension occurs gets progressively higher the further from the center the Mucus lining is displaced. The chance of host death increases with increased distension. Together, damage to the host nematode emerges as a consequence of the progression of infectious dynamics between microbial agents and with the host akin to *in vivo* observations.

While the thresholds at which intestinal swelling occurs are built into the model design, the resulting patterns of intestinal distention observed are an emergent property of the model. Experimentally, we observe that distention begins at the anterior of the intestine, and this is also observed in the model (Fig. 3). There is no difference in the rules guiding the behavior of yeast or host in the anterior; this pattern emerges from the interactions of the agents. Following the pattern-oriented modeling approach, this pattern was used to validate the model and calibrate its parameters (see Methods).

**Fig 3:**
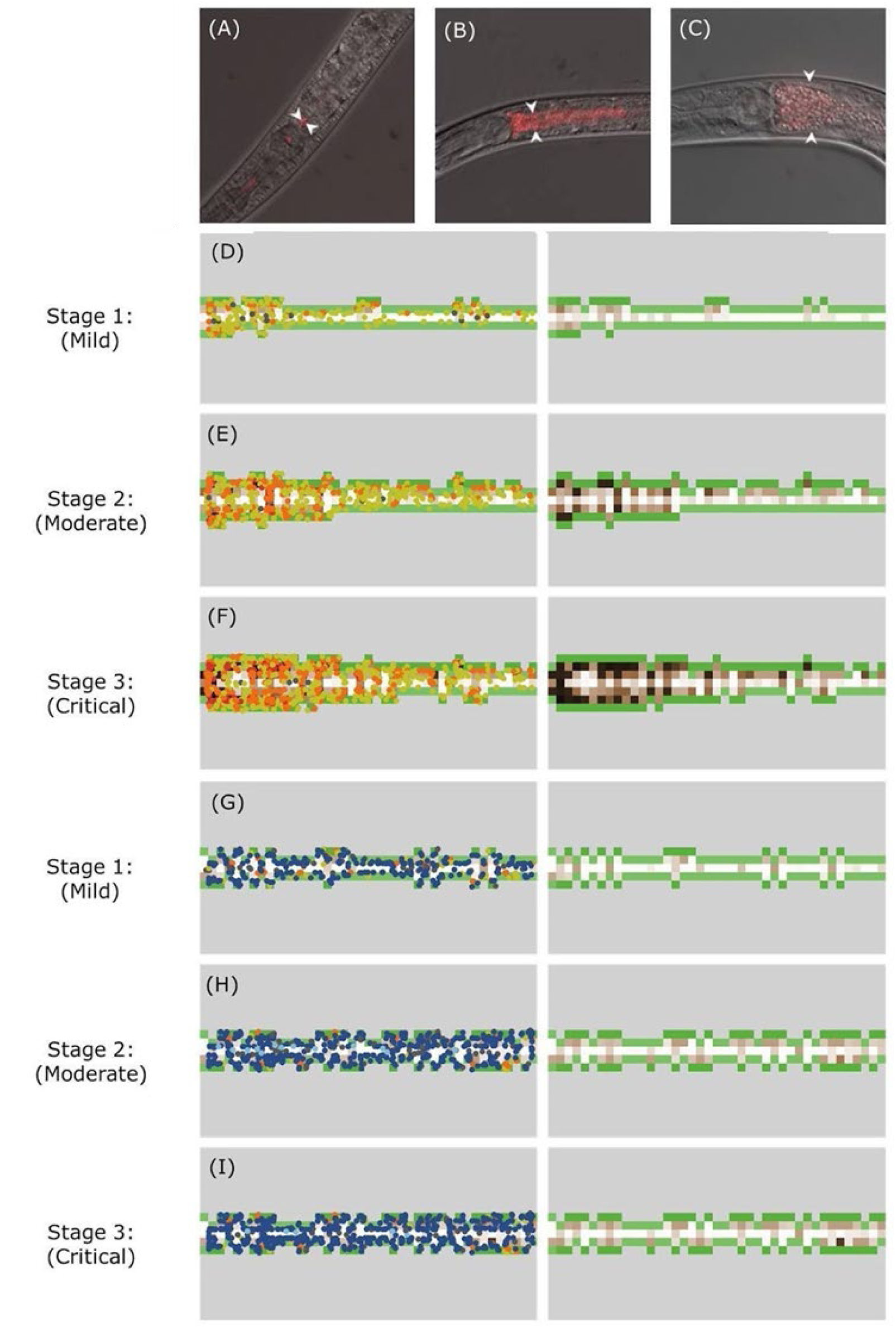
The three stages of distension. (A-C) Photomicrographs of *C. elegans* fed with pathogenic *C. albicans* labeled with mCherry for visibility. A represents the mild stage, B represents the moderate stage, and C represents the critical stage. The white arrows indicate the level of distension present in the intestine. Images were captured after two (A), three (B), and four days (C) of feeding. (D-F) Reflecting these and similar images, Enteroscape also displays these three stages in the absence of probiotic agents. D represents the mild stage, E represents the moderate stage, and F represents the critical stage. Each was captured at 1000, 2100, and 3100 ticks respectively, both with the agents shown (left panel) and hidden (right panel) for better visibility of biofilm secretion pattern. (G-I) In the presence of the probiotic yeast, pathogenic progression and severity are reduced. G-I were taken at the same respective times as D-F, after co-inoculation with pathogenic and probiotic agents. The same color scheme was used as seen in Table 1. Photomicrographs courtesy of Jain et al. [37].

### Simulation of *in vivo* infection experiments by Enteroscape reproduced observed gut distension patterns

To test the ability of Enteroscape to mimic observed patterns of infection *in vivo,* we simulated two experimental paradigms – infection with *C. albicans* alone (untreated scenario), or co-infection with *C. albicans* and probiotic yeast (treated scenario). Representative views from these scenarios at different timepoints are shown in Fig. 3.

To ensure that Enteroscape simulations were consistent with experimental results throughout infection, we broadly categorized the infection process into three stages: mild, moderate, and critical (Fig 3), and compared the appearance of these stages between experimental results and model simulations. These three stages were defined based on laboratory observations and served as benchmarks during model calibration and parameter estimation. In these simulation experiments, parameters for adhesion and de-adhesion probabilities as well as number and movement of microbes were identical for pathogenic and probiotic yeast agents.

In the mild stage in both *in vivo* experiments and simulations (Fig. 3A, 3D), little to no swelling occurred. When present, swelling typically began near the entrance of the intestine. In Enteroscape, this stage was characterized by localized “pockets” of biofilm (brown patches in Fig. 3D, right panel). The overall concentration of secreted biofilm in the model was low at this point, but early microbial colonizers and their secretions produced small, distended intestinal bulges. These bulges allowed microbes to remain adhered to a local area for longer durations than if they were attached only along the linear lumen tract. From these refuges, biofilm secretions began spreading to adjacent patches, trapping newly arriving microbes that facilitate further biofilm development.

Over time, infection progressed to the moderate stage (Fig. 3B, 3E). At this point in Enteroscape, the distinct biofilm “pockets” seen in the mild stage began to merge, forming a more continuous and cohesive biofilm structure. The characteristic intestinal bulge began to appear (Fig 3B), as these connections drove further intestinal distension. Biofilm spread from the original “pockets” and formed barriers stretching across the intestine analogous to a fishing net. The emergence of these biofilm “nets” distinguished the moderate stage from the others (Fig. 3E, right panel). As the biofilm spread horizontally and vertically, incoming microbes became caught in these “nets,” often transitioning into the Biofilm-Producing state and reinforcing the developing biofilm. This process also increased the density of microbes at the anterior of the intestine, as the biofilm barriers prevented movement towards the posterior.

In the critical stage of infection, the anterior intestinal bulge was pronounced (Fig. 3C, 3F). In Enteroscape, biofilm extended across numerous patches at high concentration (Fig. 3F, right panel). While metabolically inactive yeast occasionally appeared during the moderate stage, they primarily manifested in the critical stage due to the extent and density of biofilm accumulation. As a result, few microbes were able to move past the anterior of the intestine. Given the severity of intestinal distension, hosts typically died shortly after reaching the critical stage, both *in vivo* and in the simulation.

Simulations of the treated scenario in Enteroscape showed a marked reduction in intestinal distension (Fig 3G-I). This scenario used the same parameter values as the untreated condition, with one key difference: the co-inoculation of equal numbers of probiotic agents. The mild stage in the treated scenario resembled that of the untreated scenario (Fig. 3G), but diverged significantly in the later stages. In the presence of probiotic yeast and their secreted beneficial small molecule, swelling tended to even out across the intestinal tract. With the probiotic effect, the “pockets” of biofilm remained relatively isolated, and the adhesion of probiotic yeast agents limited the accumulation of pathogenic yeast agents. This substantially reduced biofilm production, even in what would otherwise be the critical stage in the untreated scenario. In fact, simulations of the treated scenario rarely progressed beyond the distension of late mild or early moderate stages of infection (Fig. 3H-I). Intestinal distension never reached the severity seen in the moderate or critical stages of the untreated scenario when compared to the same time periods in treated scenario simulations. Thus, the presence of the probiotic yeasts and their secreted beneficial small molecule effectively limited growth of the pathogenic yeast microbes, mirroring results reported in laboratory research [11,12].

### Simulation of *in vivo* infection experiments by Enteroscape reproduced observed survival curves

Survival analyses (Fig 4) revealed strong similarities between experimental data (panel A) and Enteroscape data (panels B and C). In both cases, infected worms that were untreated are shown in black while those treated with probiotic microbes are shown in red. Each stage of intestinal distension, as previously described, corresponds to an approximate time frame measured in ticks: the mild stage occurs around 1000 ticks, the moderate stage around 2100 ticks, and the critical stage around 3100 ticks. Although host death could occur at any stage, the majority of deaths occurred during the late moderate stage or critical stages, with a median time to death of approximately 2600 ticks across simulations using the untreated scenario (Fig. 4B-C). Hosts in the untreated scenario lived up to a maximum of 4500 ticks. When tracking the cause of death, in approximately 60% of the simulations, death resulted from infections, with the remainder due to aging.

**Fig 4:**
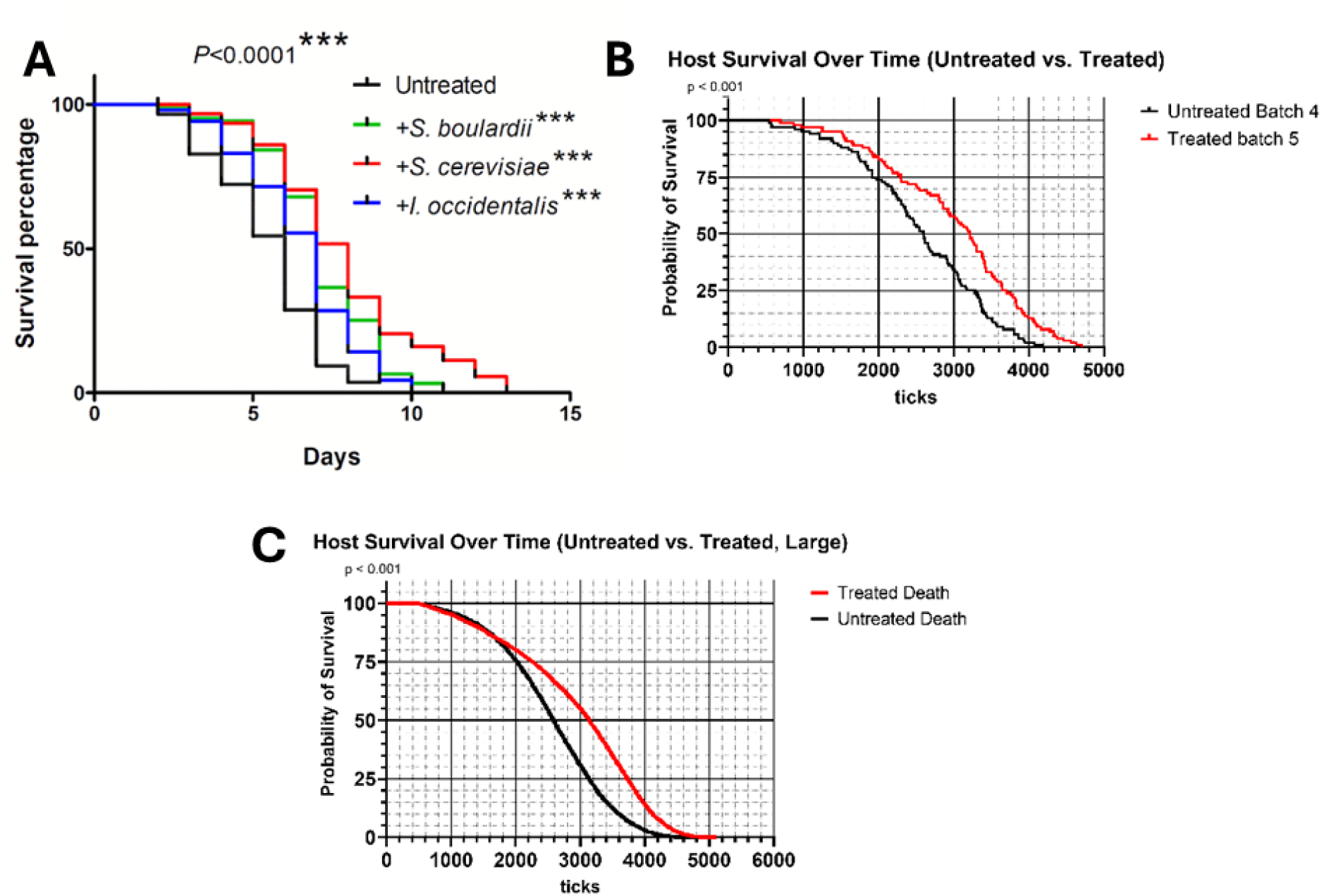
Survival curves *in vivo* and *in silico*. **(A)** Survival of *C. elegans* infected with pathogenic *Candida* across different treatments, courtesy of Kunyeit et al. (12). (n=50; log-rank test; p < 0.0001). **(B)** Survival of digital *C. elegans* simulated by Enteroscape for the untreated and treated scenarios (n = 100; log-rank test; p < 0.001). Typical untreated and treated sample curves (‘batches’) were chosen for this panel; other samples are included in the Supporting Information (S2 Appendix). (C) Large-scale survival of digital *C. elegans* simulated by Enteroscape for both the untreated and treated scenarios (n = 10,000, log-rank test; p < 0.001).

Both visually and statistically, batches with the treated scenario closely aligned with those from the untreated scenario during the early stages, before diverging around the late mild to early moderate stages (Fig. B-C). The median time to death for the treated scenario across batches was approximately 3200 ticks, with hosts typically surviving up to a maximum of 5000 ticks; this result was a statistically significant increase in lifespan compared to the untreated condition. The observed increase in both median and maximum survival times reflects a recovery in host viability relative to the untreated scenario and demonstrates a successful replication of the probiotic effect on the infectious growth of pathogenic yeast. This recovery mirrors the *in vivo* effect of probiotic yeast on survival of the host (Fig. 4A).

## Discussion

The construction of the agent-based model Enteroscape illustrates how computational sciences can enhance our understanding of biological systems. Identifying and analyzing microbe-microbe interactions alone is often challenging, and this complexity increases exponentially with the addition of each new microbial species. By leveraging both existing and ongoing research, computational approaches not only help overcome this barrier but also provide a platform for deeper investigations. Agent-based modeling is particularly valuable in this context. By representing different microbial species as distinct types of individual agents, the model enables observation of dynamics that may be difficult to capture in the laboratory. These dynamics can be readily documented through built-in digital analysis tools. Simultaneously, host-microbe interactions become more accessible when the spatial environment is treated not merely as a passive backdrop, but as an active entity with its own rules and behaviors. This approach allows the interplay between microbes, host, and environment to generate emergent phenomena that mirror laboratory observations and produce results aligned with experimental data.

A number of interesting models of the vertebrate microbiome have been developed using ABM or a combination of methods including ABM [24–26,38–40]. These models have focused on understanding how the many microbial species in the microbiome create a stable ecosystem, and how that system responds to stressors such as antibiotic treatment or different diets. Recently, the MetaBiome model used a combination of methods to simulate the heterogeneous structure and dynamics of microbial communities within the microbiome [40]. Platforms modeling microbial communities more generally and including or focusing on biofilm formation have also been developed [41–43]. All of these models include metabolic networks among microbial species as a key component of their structure. Instead, we have pursued a novel approach by establishing a simplified microbiome within the host *C. elegans* using microbes derived from the human intestine. This system allows us to explore mechanisms of infection by opportunistic pathogens such as *Candida albicans*, while uncovering microbial interactions between probiotics that can mitigate disease. In simulating this simplified experimental system, we thus chose to focus on implementing key rules governing biofilm formation and host response, rather than including metabolic networks of the microbial species.

Enteroscape includes probiotic and pathogenic yeast agents that follow known behavioral rules (secretion, adherence, etc.) and interact with each other as well as the host nematode. While we do not yet know from experimental observation that the modeled behaviors occur *in vivo*, the fact that they produce an overall phenotype in the digital host that mimics what occurs *in vivo* provides support for this hypothesis. In addition, the model makes predictions of mechanistic details that may be testable in future experiments. Enteroscape predicts not only patterns in positioning and movement of microbes along the intestine, but the formation of biofilm ‘pockets’ and ‘nets’ as well. These biofilm ‘nets’ could contribute to host mortality by inducing starvation or physiological stress. Although we haven’t documented that these nets occur *in vivo*, the model suggests a mechanism that can be experimentally tested in the future. By generating experimentally testable predictions, simulations help guide laboratory studies, which in turn inform model refinement. This iterative cycle enhances understanding of complex biological processes.

At its core, the digital model uses simplified representations to simulate the more complex coexistence observed in the original biological assay. The current version includes only two types of microbes: a pathogenic yeast and a probiotic yeast. Both share the basic functions of movement, adhesion, and de-adhesion, with differences in secretion behavior and added life cycle complexity for the pathogenic yeast agents. Likewise, the nematode intestine is represented by just three types of patches, along with two primary host behaviors: intestinal distension and death. Together, these elements reproduce the three distinct stages of infection observed in *in vivo* and *ex vivo* assays, capturing both the visual patterns of disease progression and statistical trends in survival. Running laboratory-style experiments in the model requires only basic familiarity with the platform or simply selecting from predefined experimental scenarios. The conditions and behavior of all entities can be adjusted at any time, even mid-simulation. As a foundation for experimental design and observation informed by existing research, this model holds strong potential for future applications.

There are many possible future directions for Enteroscape. One promising area of refinement lies in the secretions and interactions of the pathogenic and probiotic microbes. As described in Kunyeit et al. [12], the secretion of aromatic alcohols appear to mediate the probiotic effect of *S. cerevisiae* on the pathogenic yeasts of the *Candida* species by mitigating adhesion and biofilm formation. These effects are supported by other studies [44–46], which note that the same molecules affect hyphal growth. In both yeasts production of these aromatic alcohols is dependent on cell density and nitrogen availability, with higher production in high-density and/or nitrogen-poor conditions. Research suggests that the probiotic effect of *S. cerevisiae* via tryptophol and phenylethanol signals a nutrient-poor environment, implying a direct antagonistic relationship with *C. albicans* growth. While *C. albicans* exhibits hyphal growth at low cell densities to promote adhesion and biofilm development [45,47,48], *S. cerevisiae* undergoes pseudohyphal growth at high cell densities, possibly to aid foraging [49]. Thus, although the same molecules may signal similar environmental conditions, they elicit drastically different growth responses in each species.

Currently, Enteroscape represents this interaction in a highly simplified way, assuming a generalized “probiotic mixture” secreted solely by the probiotic agents. The exact *in vivo* secretion rates and concentrations of tryptophol and phenylethanol remain unknown. However, Enteroscape could be adapted to test hypotheses regarding more complex rules of microbial secretion and response. Simulations incorporating these hypotheses could be compared to *in vivo* phenotypes of both microbes and host; with results validated through imaging. Such expansions, alongside investigations into the biofilm ‘pockets’ and ‘nets’ mentioned previously, would not only shed light on the relationships between these organisms but strengthen the Enteroscape model further.

In summary, Enteroscape demonstrates the potential of *in silico* representations as tools for biological research. By modeling fundamental microbial behaviors, computational models can validate experimental observations and generate new hypotheses. Together, computation and laboratory experimentation expand the scope of inquiry while reinforcing core biological principles.

## Methodology

An abbreviated form of the ODD format (Objectives, Design, Details) was used here to describe the design process for Enteroscape; more technical details are available in the accompanying Supporting Information (S1 Appendix) using the full ODD protocol [34–36].

### Model Purpose and Patterns

The purpose of Enteroscape is to simulate the infectious progression of pathogenic *Candida* yeast and the probiotic effects of other yeast species on infection within the host organism *C. elegans*. By simulating this system, the model aims to enhance our understanding of host-microbe and microbe-microbe dynamics and also to support the development of potential non-pharmaceutical treatments for microbial gut infections.

Enteroscape was designed using a pattern-oriented modeling approach [22, 23, 28], inspired by experimental studies using survival assays that assess the impact of probiotic yeast species on *C. elegans* lifespan during *Candida* infection [10–12, 37]. Enteroscape’s simulations are expected to reproduce patterns observed in experimental studies, specifically the visual patterns of intestinal distension and the statistical patterns of host survival upon *Candida* infection, either alone or in combination with probiotic yeast. These patterns should emerge from a set of behavioral rules assigned to *Candida* and probiotic yeast agents, as well as the modeled *C. elegans* host response, all based on known or hypothesized mechanisms from experimental studies. Parameter calibration relied on these experimental scenarios for both visual and statistical comparisons.

### *C. elegans* Host Modeling

In Enteroscape, the *C. elegans* gastrointestinal tract is modeled using three types of patch agents: Cell (grey), Lumen (white), and Mucous patches (green) (Fig. 1). These elements simulate the anterior region of the *C. elegans* intestine, which consists of epithelial cells surrounding the intestinal lumen; these cells are each approximately 50-80 microns in length [50]. In the model, each Cell patch represents a small segment of one epithelial cell. These patches define the lumen boundary and serve as visual indicators of damage to the host: as infection progresses and the intestine distends, the number of Cell patches is reduced to reflect tissue degradation.

Lumen patches (white), represent the space within the intestine through which microbes move. These patches also monitor concentrations of microbial secretions, including beneficial small molecules and biofilm. As the simulation progresses, Lumen patches visually display these secretions using a color gradient: brown for biofilm, blue for beneficial molecules, as selected by the user (Fig. 1B). This visualization enables tracking of secretion patterns over time. Initially, microbes cannot adhere directly to Lumen patches. However, when a neighboring pathogenic yeast agent adheres to a patch and begins secreting biofilm, that biofilm spreads to adjacent Lumen patches. Once biofilm is present, the likelihood of microbial adhesion increases with its concentration. *In vivo,* the anterior region of the *C. elegans* intestine has a more complex morphology than the rest of the intestine [51]; for simplicity, Enteroscape represents this area with an enlarged lumen as compared to the rest of the intestine.

Mucous patches (green) simulate the mucosal lining of the nematode intestine. In living nematodes, this lining has microvilli and a dense layer of proteins and extracellular matrix called the glycocalyx, which serves as a protective barrier for epithelial cells and also aids in digesting food [51]. In Enteroscape, Mucous patches are considered maximally adherent by default, due to the glycocalyx. Microbes initially adhere to these Mucous patches; however, pathogenic yeast agents do not deposit biofilm onto Mucous patches. Instead, secreted biofilm is redirected to surrounding Lumen patches, promoting further biofilm accumulation and infection progression.

### Microbial Modeling – Life Cycle

Both microbial agent types use a modified Brownian motion to navigate through the intestine. In C. *elegans*, intestinal flow is directed posteriorly due to pharyngeal pumping and the defecation motor program, which push food/microbes towards the posterior end of the intestine [52,53]. In Enteroscape, this behavior is modeled by constraining the heading (direction of motion) of microbial agents so they generally move towards the posterior.

Both agent types can exist in one of the two initial states: Planktonic (referred to in the model as ‘Unstuck’) or Adherent (‘Stuck’). Agents may transition between these states at each time step (tick) of the simulation (Fig. 2). In the Planktonic state, microbial agents move freely through the intestinal Lumen patches and slowly through Mucous patches. When a pathogenic microbe agent encounters a Mucous patch or a Lumen patch containing secreted biofilm, it has a chance to transition to the Adherent state. This probability is governed by user-adjustable global variables called ‘Yeast-adhesion-prob.’ The likelihood of adhesion increases with the local concentration of biofilm on a Lumen patch. In the Adherent state, the microbe agent is immobilized. It can return to the Planktonic state with a probability set by another global variable, ‘Yeast-deadhesion-prob.’ As with adhesion, the probability of de-adhesion decreases as biofilm concentration on the patch increases. Probiotic microbial agents follow the same state transition rules, but use separate global variables for adhesion and de-adhesion: ‘Beneficial-Microbe-adhesion-prob’ and ‘Beneficial-Microbe-deadhesion-prob’ respectively.

The key behavioral differences between probiotic and pathogenic yeast microbial agents lies in their secretion. In alignment with both *ex vivo* and *in vivo* observations, probiotic yeast agents continuously secrete a small beneficial molecule regardless of whether they are in the Adherent or Planktonic state. This beneficial small-molecule represents the secretion of metabolites that inhibit pathogenic yeast adhesion as identified in previous studies [12]. Within the model, this beneficial molecule functionally competes with the biofilm secreted by pathogenic yeast agents. Both secretions are modeled as patch variables that increase when a microbe secretes onto a patch. While biofilm increases the probability of adhesion and decreases the probability of de-adhesion, the beneficial small-molecule has the opposite effect: it reduces adhesion and promotes de-adhesion. The rate of degradation can be controlled by the user by the Beneficial-Microbe-molecule-degradation-rate slider on the Enteroscape model interface.

If a sufficient number of pathogenic yeast agents remain adherent in a region for an extended period, they may transition to a third pathogen-specific state: Biofilm-Producing. These agents remain adhered but begin secreting biofilm onto their local patch every tick. If the patch reaches its maximum biofilm capacity, the excess biofilm spreads to the adjacent patches that have not yet reached saturation. Biofilm-Producing pathogenic yeast agents cannot de-adhere and return to the Planktonic state, as they have now actively integrated into the biofilm structure. When a Biofilm-Producing pathogenic yeast agent becomes surrounded by high concentrations of biofilm, it transitions into a fourth state and final state: Metabolically Inactive. This state reflects the reduced metabolic activity observed in microbes located in the dense, mature center of biofilms [14,54]. While biofilms protect embedded microbes from drugs and other stressors, they also limit nutrient availability. As a result deeply embedded microbes downregulate nonessential functions to survive. In this model, Metabolically Inactive pathogenic yeast agents cease biofilm secretion and are considered fully incorporated into the biofilm. Their presence signals advanced infection progression, as only regions with high biofilm concentration allow agents to reach this terminal state.

### Microbial Interactions -- Spatial Mechanics

*C. elegans* intestinal lumen has limited physical space, and Enteroscape reflects this by restricting the number of microbes that can occupy a single patch. In addition to microbes de-adhering on their own, laboratory observations suggest that external forces like the movement of neighboring microbes can also dislodge them. Enteroscape incorporates this phenomenon by allowing microbial agents that attempt to move into an already fully occupied patch a chance to dislodge some of the current residents, or be displaced themselves. This is implemented by identifying patches where the number of microbial agents reaches a user-defined global threshold. When such overcrowding occurs, a subset of microbial agents is randomly selected for displacement until the patch falls below the threshold. This process can result in several outcomes: the incoming agent may be repelled (‘bounce off’), one or more of the existing adherent agents may be displaced, or both may be displaced simultaneously. Pathogenic yeast agents in the Biofilm-Producing and Metabolically Inactive states are exempt from this displacement as they are considered to be fully integrated into the surrounding biofilm, However, Adherent agents of either microbial type can be dislodged and revert to the Planktonic state.

These spatial dynamics influence the distribution and movement of microbes along the intestinal tract and enhance the biological realism of Enteroscape’s simulation of microbial interactions.

### Host-Microbe Interactions -- Intestinal Swelling and Distension

In the context of host-microbe interactions, the primary outcome of pathogenic yeast overgrowth is damage to the host tissue. Consistent with *in vivo* observations, Enteroscape represents this damage through intestinal distension. As described earlier, Mucous patches monitor the key conditions that can trigger swelling: the number of microbial agents present on the patch and the average concentration of biofilm on neighboring patches. Unlike the displacement mechanism, the distension mechanism accounts for microbes in any state. The distension mechanism also accounts for surrounding patches, rather than each patch individually as the displacement mechanism does. When these conditions are met, swelling is visually represented by the outward displacement of Mucous patches from the intestinal center line. This is achieved through a transformation of patch types: the affected Mucous patch becomes a Lumen patch (effectively expanding the Lumen), while the adjacent Cell patches are converted to new Mucous patches.

The resident microbe agents move vertically with this expansion, with those originally on the Mucous patch repositioned onto the newly formed Mucous patch above. Because more severe distension reflects greater pathogenic activity, the criteria for continued swelling beyond the initial displacement becomes increasingly stringent. Higher levels of yeast presence and biofilm concentration are required to drive further expansion. The model defines the severity of intestinal distension by measuring the absolute distance between Mucous patches and the intestinal center.

### Determination of Host Death

In Enteroscape, the probability of host death increases as a function of both host age and severity of distention of the intestine. Using existing mortality studies in *C. elegans* [55–62], the model incorporates an age-based mortality component defined by a user-defined baseline percentage. This probability increases incrementally every time the model checks for host death due to age or other infection-independent factors. As a result, microbial agents display a characteristic exponential decline in host survival, reflecting natural survival/aging patterns in nematodes.

The infection-related mortality component is driven by the extent of intestinal distention. This is calculated as a weighted average of the distance each mucous patch has shifted from the intestinal midline. A sigmoid function is used to convert this average distance into a probability of death. Mucous patches at their original locations contribute negligibly to the death probability, while those further from the midline contribute increasingly: with the likelihood approaching nearly one hundred percent at a displacement of 5 patches.

To avoid the need for unrealistically small per-tick death probabilities, Enteroscape does not evaluate host death continuously. Instead, it checks both age-related and infection-related mortality conditions at pseudo-random intervals. This approach maintains a realistic frequency of checks while ensuring that the two mechanisms, age and infection, remain independent and non-arbitrary in their contribution to host death within the model.

### Parameter Calibration

Enteroscape uses the pattern-oriented modeling approach, which reduces uncertainty in the selection of parameter values through two key mechanisms [22]. First, because the model is built upon realistic representation of underlying biological processes, its outputs are generally less sensitive to uncertainty in individual parameter values [63]. Second, parameters are tuned based on their ability to reproduce a range of observed patterns, rather than a single outcome; this process is known as “inverse modeling” [64]. In our simulation, these patterns include both visual infection dynamics at different stages of disease progression and quantitative outputs such as survival curve trajectories and statistical differences between treatment conditions. Importantly, these emergent patterns arise from the fundamental behavioral rules embedded in the model; they are not explicitly coded into or defined by the model design.

### Infection Scenarios

As with its construction, Enteroscape’s calibration is grounded in empirical data from laboratory experiments and previous studies. To support comparison with *in vivo* observations, the model includes two preprogrammed infection scenarios derived from previous research: an untreated scenario and a treated (co-inoculated) scenario. The untreated scenario simulates a control condition in which the host is exposed only to pathogenic yeasts. In contrast, the treated scenario simulates a condition in which both pathogenic and probiotic yeasts are introduced simultaneously (co-inoculation scenario). Although earlier research also explored pre- and post-inoculation treatment strategies [11,12] this study focuses on the co-inoculation condition, as it demonstrated the most robust effect *in vivo*.

### Visual calibration

Visual calibration of Enteroscape involved comparing simulation outputs to *in vivo* images of infection progression in nematodes that were fed fluorescently labeled *Candida albicans* in the laboratory. As discussed previously, achieving similar visual patterns of infection in the simulation suggests that the model captures the underlying biological processes with reasonable accuracy. Importantly, this comparison focused not only on the final stages of infection, but also the intermediate stages of disease progression *in vivo*. The ability of Enteroscape to reproduce these dynamic visual patterns provided evidence that the model accurately mirrors key aspects of real biological system behavior and infection dynamics.

### Statistical calibration

To further evaluate the model’s accuracy, we conducted statistical calibration through survival analysis. Using survival data and analysis from previously published studies, we compared lifespan trends from Enteroscape simulations to those observed in experiments *in vivo*. Specifically, we analyzed both untreated and treated (co-inoculated) scenarios using batch runs generated with NetLogo’s BehaviorSpace tool. Each batch consisted of approximately 100 simulated replicates, roughly twice the sample size (n=50) used in the reference studies [11,12]. Simulation outputs were used to construct Kaplan-Meier survival curves, enabling direct comparison to *in vivo* survival data. Statistical tests, including the log-rank and Gehan-Breslow-Wilcoxon methods were used to assess significance in lifespan differences between treatment conditions. The ability of Enteroscape to reproduce not only the shape of the survival curves but also statistically significant differences between treatment groups supports its validity as a model of host-microbe interactions.

### Model Parameter Choices

Finalizing the default parameter values for Enteroscape required extensive iterative testing to ensure that simulations reliably produced the expected emergent patterns. The calibration process began with repeated simulation runs to identify approximate acceptable ranges for key parameters. Using these baseline estimates, we performed targeted single-variable sweeps for several visible parameters including: Beneficial-Microbe-molecule-secretion-rate, Beneficial-Microbe-molecule-degradation-rate, Yeast-biofilm-secretion-rate, and Worm-Death-Probability. These sweeps helped tune the system to replicate the infection dynamics and survival outcomes observed in the *in vivo* scenarios. Other parameter pairs such as Yeast-adhesion-prob and Yeast-deadhesion-prob, or their counterparts for probiotic yeast Beneficial-Microbe-adhesion-prob and Beneficial-Microbe-deadhesion-prob, were tested in tandem, due to their interdependent roles in microbial adhesion dynamics. These paired sweeps explored all possible value combinations within identified ranges to capture their joint effects on infection progression. In addition to parameters visible to the user, several internal system variables not directly adjustable through the user interface were also tested. These included: g-increasing-chance and g-natural-check which control how frequently the model checks for host death; the parameters for the sigmoidal function used to calculate the probability of death due to intestinal distension; and g-Biofilm-Swell-Thresh, the base threshold of biofilm concentration required to trigger distension of the host intestinal.

Each parameter sweep consisted of 100 simulation replicates per value, following the same batch-run methodology used in survival analysis. Outcomes were assessed using both statistical survival data and visual inspection of infection progression at each of the three defined stages. This dual calibration approach ensured that the chosen parameters produced biologically plausible dynamics without bias towards either statistical or visual outcomes. The values identified by these parameter sweeps were used to construct the default settings for the untreated and treated (co-inoculated) infection scenarios, and to fine tune the model’s overall behavior. The result is a validated, well-calibrated version of Enteroscape capable of capturing both mechanistic and emergent aspects of microbial interactions in the *C. elegans* intestine.

## Acknowledgements

The authors would like to recognize several individuals for their assistance with the development of this project. Sharada Vishwanath, Nicholas Tourtillott, Daire Daley, and Monet Norales helped design the initial mechanics and concept of the model from which Enteroscape was developed. In addition, Dr. Romina D’Almeida and Dr. Lohith Kunyeit from the Rao Lab helped contribute imaging and details necessary for the *in vivo* comparison and validation of model results.

## Supporting Information Captions

**S1 Appendix. Enteroscape ODD protocol (Objectives, Design, Details).** This document contains the full technical description of the Enteroscape model.

**S2 Appendix. Additional survival curve data generated by Enteroscape**.

## Data Availability Statement

The Netlogo file, source code, and optimization data used to produce the analysis and results in this manuscript are available in this GitHub repository: https://github.com/daviddatt/Enteroscape.

